# Inositol depletion regulates phospholipid metabolism and activates stress signaling in HEK293T cells

**DOI:** 10.1101/2022.02.21.481362

**Authors:** Mahmoud Suliman, Kendall C. Case, Michael W. Schmidtke, Pablo Lazcano, Chisom J. Onu, Miriam L. Greenberg

## Abstract

Inositol plays a significant role in cellular function and signaling. Studies in yeast have demonstrated an “inositol-less death” phenotype, suggesting that inositol is an essential metabolite. In yeast, inositol synthesis is highly regulated, and inositol levels have been shown to be a major metabolic regulator, with its abundance affecting the expression of hundreds of genes. Abnormalities in inositol metabolism have been associated with several human disorders. Despite its importance, very little is known about the regulation of inositol synthesis and the pathways regulated by inositol in human cells. The current study aimed to address this knowledge gap. Knockout of ISYNA1 (encoding *myo*-inositol-3-P synthase 1) in HEK293T cells generated a human cell line that is deficient in *de novo* inositol synthesis. ISYNA1-KO cells exhibited inositol-less death when deprived of inositol. Lipidomic analysis identified inositol depletion as a global regulator of phospholipid levels in human cells, including downregulation of phosphatidylinositol (PI) and upregulation of the phosphatidylglycerol (PG)/cardiolipin (CL) branch of phospholipid metabolism. RNA-Seq analysis revealed that inositol depletion induced substantial changes in the expression of genes involved in cell signaling, including extracellular signal-regulated kinase (ERK), and genes controlling amino acid transport and protein processing in the endoplasmic reticulum (ER). This study provides the first in-depth characterization of the effects of inositol depletion on phospholipid metabolism and gene expression in human cells, establishing an essential role for inositol in maintaining cell viability and regulating cell signaling and metabolism.

## 1. INTRODUCTION

Inositol is an essential carbocyclic sugar that plays a vital role in cell signaling, intracellular trafficking, and osmoregulation. Inositol can be obtained from extracellular sources through the action of inositol transporters or synthesized *de novo* from glucose [1–3]. The first and rate-limiting step of inositol synthesis is the conversion of glucose-6-phosphate to inositol-3-phosphate (I3P) by the enzyme *myo*-inositol-3-phosphate synthase (MIPS) [3], which was first identified and purified from yeast and mapped to the *INO1* gene [4]. Inositol synthesis in yeast is highly regulated. Yeast *INO1* is transcriptionally regulated by the Henry regulatory circuit, including activator protein complex Ino2-Ino4 and repressor Opi1, which bind to the upstream activating sequence UAS_INO_ [5–7]. This regulation is dependent on the presence of extracellular inositol, as *INO1* expression is almost completely repressed when yeast cultures are supplemented with inositol [3]. The second step of inositol synthesis is the dephosphorylation of I3P by inositol monophosphatases to form free inositol [8]. Inositol can be subsequently combined with CDP-diacylglycerol (CDP-DAG) by phosphatidylinositol synthase to form the phospholipid phosphatidylinositol [5]. Additional phosphorylation of the inositol head group produces a range of phosphoinositide species [3]. Phosphoinositides are critical for a myriad of cellular processes, including exo- and endocytosis, cytoskeleton dynamics, protein trafficking, membrane homeostasis, and cell survival and motility [9]. Inositol and I3P are the precursors of the water-soluble inositol phosphates and inositol pyrophosphates [3, 10, 11, which play a significant role in cell signaling and proliferation, egg fertilization, neuronal development, and regulation of energy metabolism [3, 10]. Inositol phosphate and pyrophosphate synthesis utilizes inositol derived from extracellular sources preferentially over inositol synthesized *de novo* [12]. Surprisingly, inositol phosphates can be synthesized from glucose even in the absence of MIPS, albeit to a lesser extent. This finding suggests the existence of an additional less-utilized pathway for generating inositol-containing compounds from glucose [12], and illustrates that much still remains to be elucidated regarding the critically important and highly regulated metabolism of inositol.

Inositol has been shown to be essential for cell growth and viability in all eukaryotes studied to date. Inositol-less death has been extensively characterized in budding yeast and other fungi [13–15]. *Schizosaccharomyces pombe*, a natural inositol auxotroph lacking functional MIPS, dies without supplementation of exogenous inositol in the growth medium [16]. Similarly, the *Saccharomyces cerevisiae ino1*Δ strain is also an inositol auxotroph, as it lacks the gene encoding MIPS [17]. Cell division is arrested within two hours of transferring these cells to inositol-free medium, with cell death commencing after four hours [17]. This is not surprising, given that inositol robustly affects gene transcription in yeast, regulating hundreds of genes and pathways related to phospholipid metabolism, the unfolded protein response, amino acid metabolism, and ribosome biogenesis [18–20]. Inositol is also a critical supplement used in mammalian cell culture, and inositol deprivation has been shown to induce cell death in several mammalian cell lines [21–23]. Recent studies demonstrated that disruption of both *de novo* inositol synthesis and inositol transport leads to inositol auxotrophy in leukemia cells [21], and knockout of MIPS from human colon cancer cells results in a severe reduction in cell proliferation when grown in inositol-free medium [12]. Furthermore, an inositol-deficient diet led to reduced growth, and impaired the intestine function of grass carp [24]. Overall, these studies suggest that the requirement for inositol is conserved across all eukaryotes.

Abnormalities in inositol metabolism have been associated with several human disorders such as bipolar disorder, Alzheimer’s disease, polycystic ovary syndrome, and cancer [25, 26]. *Myo*-inositol, the most abundant isoform (herein referred to as inositol), can be interconverted into other isomers. These include D-*chiro*-inositol and *scyllo*-inositol, which have also been shown to play an essential role in human health [27]. Despite the vital importance of inositol, very little is known about its function and regulation in humans.

The current study aimed to address this knowledge gap by rigorously characterizing the consequences of inositol depletion in human cells. Using CRISPR technology, a human embryonic kidney (HEK293T) cell line incapable of *de novo* inositol synthesis was generated by disrupting the inositol-3-P synthase 1 (ISYNA1) gene, the mammalian homolog of yeast *INO1*. Using this ISYNA1-KO cell line in combination with a uniquely derived inositol-free culture medium permitted the first in-depth characterization of inositol depletion in human cells. This study demonstrates that inositol depletion induces profound changes to the cellular phospholipid profile, most notably downregulation of phosphatidylinositol (PI) and upregulation of the mitochondrial phosphatidylglycerol (PG)/cardiolipin (CL) branch of phospholipid metabolism. Furthermore, transcriptomic analysis identified ~300 genes that are responsive to inositol depletion, including robust activation of the ERK and ATF4 signaling pathways and upregulation of autophagy. These findings underscore the vital importance of inositol regulation in human cells and may provide valuable insights into the mechanistic effects of inositol-depleting drugs.

## 2. MATERIALS AND METHODS

### 2.1. Cell line and growth conditions

HEK293T cells were maintained in DMEM (GIBCO), 10% fetal bovine serum (FBS) (ATLANTA), 1% pen-strep. To characterize the effect of inositol on cell growth and physiology, cells were cultured in inositol-free DMEM (MP BIO) supplemented with 4% glutamine and lab-dialyzed FBS (Cytiva Life Sciences, Marlborough, MA). Dialysis was performed against deionized water using Spectra/Por 3 dialysis membrane tubing (Repligen) with a MWCO of 3.5 kDa for 48 hours. The water in the container was replaced four times per day.

The yeast strain *ino1*Δ used in this study belongs to the *Saccharomyces cerevisiae* BY4741 background and was obtained from Invitrogen. Cells were maintained on YPD plates (2% bactopeptone, 1% yeast extract, 2% glucose, 2% agar) and cultured in synthetic complete (SCD) medium, which contains glucose (20 g/L), ammonium sulfate (2 g/L), yeast nitrogen base (0.69 g/L), adenine (20.25 mg/L), arginine (20 mg/L), histidine (10 mg/L), leucine (60 mg/L), lysine (20 mg/L), methionine (20 mg/L), threonine (300 mg/L), tryptophan (20 mg/L), and uracil (40 mg/L) and inositol-free vitamin mix. To maintain the *ino1*Δ inositol auxotrophic strain, SCD medium was supplemented with 75 μM inositol. Cells were incubated at 30°C, with shaking for liquid cultures.

### 2.2. Construction of the ISYNA1-KO cell line using CRISPR technology

The HEK293T ISYNA1 knockout cell line was constructed using a double nickase system (Santa Cruz Biotechnology sc-404785-NIC) that contains a pair of plasmids each expressing a D10A mutated Cas9 nuclease and 20 nucleotide guide RNA (gRNA) designed to target exon 1 of ISYNA1. To confirm successful gene disruption, Western blot analysis was performed using antibodies against the ISYNA1 translated protein, MIPS. To obtain the DNA sequence of the individual alleles, DNA was extracted from cells, and the ISYNA1 genomic region was amplified by PCR (forward primer: ATGAATTCTCGAACCGGGTCTCCAG, reverse primer: TAGCGGCCGCCAGGACAAACGCAGTCGAT). Amplicons were cloned with EcoRI and NotI into the pGKB vector and transformed into *E. coli* to isolate the different alleles. Eight transformants of pGKB vectors containing one of the two alleles were direct colony sequenced by Genewiz using a primer sitting on the promoter upstream of the multiple cloning site (AATAATAGCGGGCGGACGCA).

Off-target analysis for the ISYNA1-KO cell line was performed by first identifying the top 10 predicted off-target loci using the CCTop CRISPR/Cas9 target predictor tool (https://crispr.cos.uni-heidelberg.de/). This software compares the genomic ISYNA1 sequence targeted by the CRISPR/Cas9 gRNA to the entire human genome (mapped to reference genome hg19) to identify additional loci that have sequence similarity to the target locus. Next, whole-genome sequencing and differential variant analysis (Psomagen Inc., Rockville, MD) were performed on both the WT and ISYNA1-KO cell lines to compare the sequences at each potential off-target locus (Table S4). This analysis confirmed the targeted disruption of exon 1 in the ISYNA1 gene and revealed that none of the top 10 predicted off-target sites were altered in the ISYNA1-KO cell line relative to the WT cells from which they were derived.

### 2.3. Construction of ISYNA1 expression plasmids

Human ISYNA1 isoform 2 (UniProt classification) was cloned into a pcDNA4/TO mammalian expression vector (Thermo Fisher Scientific). The cDNA was amplified by PCR (forward primer: GTTTAAACTTAAGCTTATGGAGGCCGCCGC, reverse primer: AAACGGGCCCTCTAGATCAGGTGGTGGGCATTGG) from a previously cloned plasmid harboring the ISYNA1 gene [28]. Amplicons were cloned into the pcDNA4/TO vector with XbaI and HindIII. Plasmids were purified from *E. coli* using the NucleoSpin Plasmid miniprep kit (Takara 740588). Transformants were sequenced to confirm successful cloning. As a control, we used the same vector expressing lacZ protein.

### 2.4. Cell viability assay

Cell viability assays were performed using Cell Count Kit 8 (CCK-8; GLPBIO, USA) according to the manufacturer’s protocol. CCK-8 utilizes a highly water-soluble tetrazolium salt, which produces a water-soluble formazan dye upon reduction by dehydrogenases in cells to give an orange-colored product. The amount of the formazan dye generated is directly proportional to the number of living cells. Cells were cultured in 96-well plates (~700 or ~1000 cells/well) in 150 uL of DMEM supplemented with 10% dialyzed FBS. For each experimental condition, cells were cultured in 6-8 triplicates, and cell viability was measured at the indicated days using one of the triplicates. 15 uL of assay reagent was added to each well, and A_450_ was measured after 3 hours. Additionally, a standard curve of cell number vs. A_450_ was generated and used to convert each A_450_ reading into a viable cell number. For cell viability experiments with the ISYNA1 expression plasmid, cells were cultured in a T-25 flask and transfected with 250 ng of the ISYNA1 expression plasmid or control plasmid expressing LacZ protein 48 hours before subculturing into 96-well plates and performing the cell viability assay.

### 2.5. Immunoblotting

Cell lysates were extracted using a radioimmunoprecipitation assay (RIPA) buffer (Santa Cruz Biotechnology). Protein concentration was determined using the Pierce BCA Protein Assay (Thermo Fisher Scientific). 30 μg of protein mixed with loading buffer (62.5 mM Tris pH 6.8, 10% (v/v) glycerol, 2% (w/v) SDS, 0.01% (w/v) bromophenol blue) was run on SDS-PAGE gels (15% for LC3B analysis and 12% for the other proteins analyzed) then transferred to a polyvinylidene difluoride (PVDF) membrane. Protein detection was done using primary antibodies against ISYNA1 (H-300, rabbit, Santa Cruz Biotechnology), β-actin (C-4, mouse, Santa Cruz Biotechnology), P44/42 Erk1/2 (#9102, Cell signaling), Phospho-p44/42 MAPK (Erk1/2) (#4370, Cell signaling), ATF-4 (#11815, Cell signaling), LC3B (#NB600-1384, Novus Biologicals), and Cyclophilin B (#43603, Cell signaling). For secondary antibodies, the corresponding europium-labeled ScanLater Western Blot Assay Kit (Molecular Devises, USA) was used, and membranes were imaged using a SpectraMax i3x plate reader with the ScanLater module (Molecular Devices, San Jose, CA). ImageJ was used to quantify the intensity of bands, which were normalized to housekeeping proteins or total protein stain. Values are shown as fold change relative to control.

Yeast cell lysates were extracted using a boiling technique [29]. Briefly, 3 mL of yeast cultures in mid-log growth phase were directly incubated at 100°C for 5 min and dead cells collected by centrifugation at 3000 x *g* for 5 min. The pellet was incubated with 150 μL TE buffer (10 mM Tris-HCl, 1 mM EDTA, pH 7.5) and 150 μL 0.2 M NaOH for 5 min and transferred to a microcentrifuge tube. Cells were then collected by centrifugation at 11,000 x *g* for 1 min, resuspended in loading buffer at 30 μL/1.0 OD_600_ x mL cells collected, then incubated at 100°C for 5 min. 10 uL of these normalized extracts were run on an 8-16% SDS-PAGE gel for 2 hrs at 120V then transferred to a PVDF membrane at 30V for 18 hrs at 4°C. Total protein staining of membrane was done using an R-PROB staining kit from Sigma. Primary antibodies were incubated overnight at 4°C and secondaries for 1 hr at RT. Protein detection was done using a custom antibody against MIPS (1:10,000) and anti-actin (1:10,000; Invitrogen MA1-744). The custom MIPS antibodies were produced by YenZym Antibodies, LLC by raising rabbits against the synthesized peptide SYNHLGNNDGYNLSAPKQFRSKE-Ahx-C targeting amino acids 348-370 of the *Saccharomyces cerevisiae* MIPS protein. Antibodies were purified from serum using the peptide antigen.

### 2.6. Lipidomic analysis using HPLC-MS

Lipidomic analysis was performed by the Core Facility Metabolomics of the Amsterdam UMC as described previously [30]. The HPLC system consisted of an Ultimate 3000 binary HPLC pump, a vacuum degasser, a column temperature controller, and an autosampler (Thermo Scientific, Waltham, MA, USA). The column temperature was maintained at 25°C. The lipid extract was injected onto a “normal phase column” LiChrospher 2×250-mm silica-60 column, 5 μm particle diameter (Merck, Darmstadt, Germany) and a “reverse phase column” Acquity UPLC HSS T3, 1.8 μm particle diameter (Waters, Milford Massachusetts, USA). A Q Exactive Plus Orbitrap (Thermo Scientific) mass spectrometer was used in the negative and positive electrospray ionization mode. In both ionization modes, mass spectra of the lipid species were obtained by continuous scanning from m/z 150 to 2000 with a resolution of 280,000 full widths at half-maximum (FWHM). The RAW HPLC-MS data files were converted to mzXML using MSconvert [31] in centroided mode. Peak finding and peak group finding were done using the R package XCMS, with minor modifications to some functions for a better representation of the Q Exactive data. Annotation of the peaks was done based on an in-house database containing all possible (phospho)lipid species. Each combination of column (normal-phase or reverse-phase) and scan mode (positive or negative) was processed separately; after normalization, separate peak group lists were combined into two resulting lists, which were used for statistical analysis.

### 2.7. RNA-Seq and gene set enrichment analysis

Total RNA was extracted from cells using TRIzol Reagent (Invitrogen) following the manufacturer’s instructions. Expression analysis was conducted in collaboration with the Wayne State University Genome Sciences Core as described previously [32]. Four biological replicates were used for each condition. QuantSeq 3’ mRNA-Seq Library Prep Kit FWD for Illumina (Lexogen) was used to generate libraries of sequences close to the 3’ end of polyadenylated RNA from 15 ng of total RNA in 5 μL of nuclease-free water following a low-input protocol. Library aliquots were assessed using the Qubit 1X dsDNA HS Assay kit (Invitrogen). Barcoded libraries were normalized to 2 nM before sequencing at 300 pM on one lane of a NovaSeq 6000 SP flow cell. After de-multiplexing with Illumina’s CASAVA 1.8.2 software, the 75 bp reads were aligned to the human genome (Build hg38) with STAR_2.4 [33] and tabulated for each gene region [34]. Differential gene expression analysis was used to compare transcriptome changes between conditions using edgeR v.3.22.3 [35], and transcripts were defined as significantly differentially expressed at a false discovery rate (FDR) ≤ 0.05.

Gene enrichment analysis of significantly differentially expressed genes was performed using ShinyGO (v0.741). The enrichment was performed for (A) GO molecular functions (B) Kyoto Encyclopedia of Genes and Genomes (KEGG) pathways.

### 2.8. Heatmaps

Heatmaps were generated using pheatmap v1.0.12 in R [36, 37]. For the lipidomic data, the heatmaps were generated using row Z-scores of relative abundance units for each lipid species across replicates and conditions. For the RNA-Seq data, the heatmaps were generated using row Z-scores of counts representing the number of reads mapped for each gene across replicates and conditions. The formula for calculating a Z-score is **z = (x-μ)/σ**, where x is the raw score, μ is the mean, and σ is the standard deviation.

### 2.9. Statistical analysis

The data are shown as mean ± standard deviation. Results are from at least three biological replicates. Proper statistical tests were performed (e.g., Student’s t-test) to conclude statistical significance (*p* ≤ 0.05).

## 3. RESULTS

### 3.1. Construction of a HEK293T cell line deficient in de novo inositol synthesis

To study the effects of inositol deprivation in human cells, an ISYNA1-KO cell line was constructed in HEK293T cells. The ISYNA1 gene encodes the enzyme MIPS, which mediates the first and rate-limiting step in *de novo* inositol synthesis. HEK293T is a well-characterized cell line that has been used extensively for expression and biochemical studies [38]. To generate this model, cells were transfected with CRISPR/Cas9 plasmids designed to disrupt exon 1 of ISYNA1. A monoclonal lineage of mutated cells was isolated, and Western blot analysis was performed to confirm the absence of MIPS protein (Fig. 1a). To further authenticate this clone, the two ISYNA1 alleles were sequenced, revealing frameshift mutations at the targeted locus. These mutations not only changed the amino acid sequence relative to WT MIPS, but also caused truncation of the protein by insertion of a premature stop codon (Fig. 1b). To ensure the absence of off-target mutations, whole-genome sequencing of isogenic WT and ISYNA1-KO cells was conducted, and a differential variant analysis was used to confirm the absence of off-target mutations at any of the top 10 predicted off-target loci (Table S4). This novel cell line, lacking MIPS and therefore completely deficient in inositol synthesis, was utilized in the following experiments to determine the cellular consequences of inositol deprivation.

**Figure 1.**
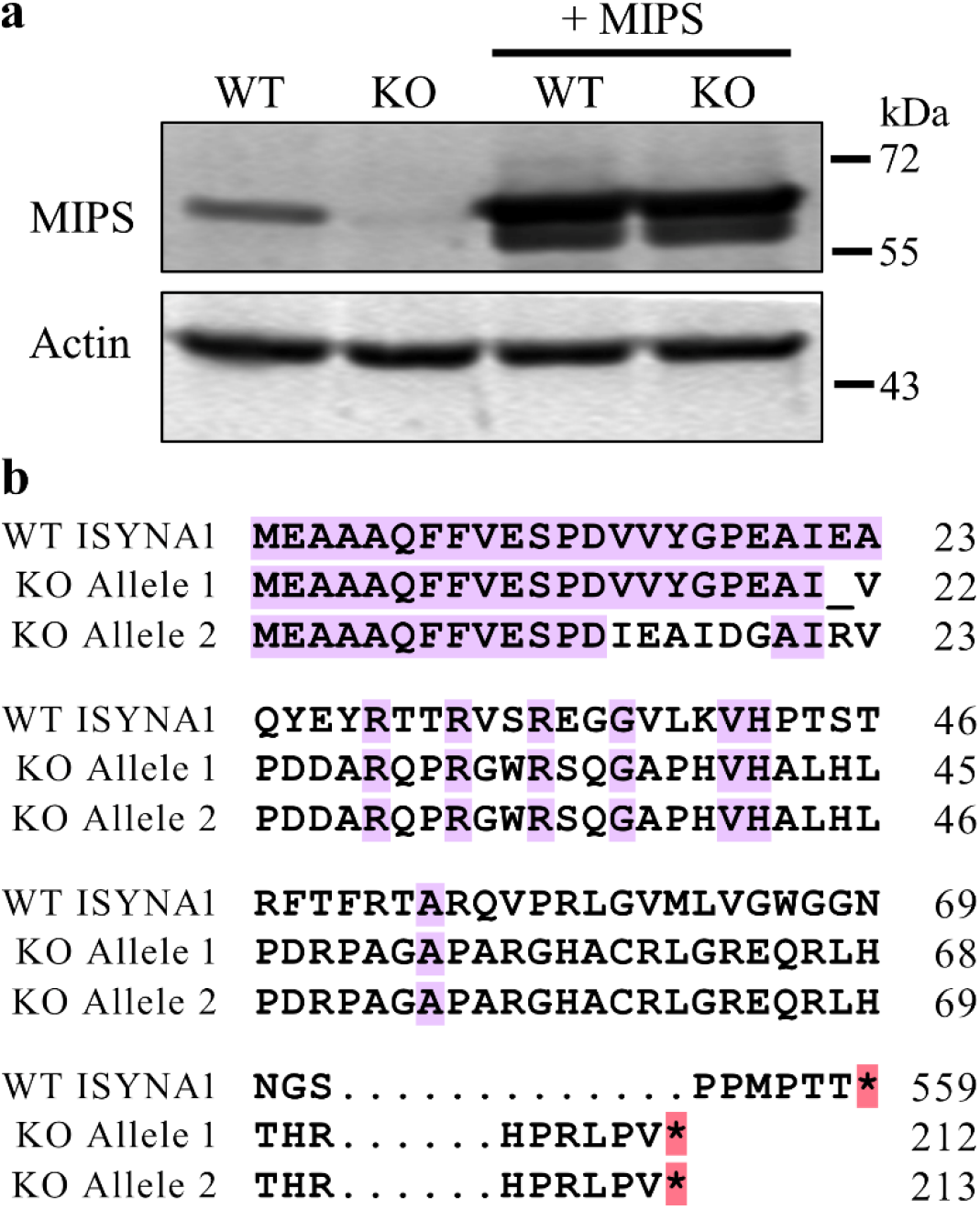
Confirmation of the HEK293T ISYNA1-KO cell line and ISYNA1 expression plasmid. A HEK293T ISYNA1 knockout cell line was constructed using CRISPR/Cas9 plasmids designed to disrupt exon 1 of ISYNA1. Where indicated, WT and KO cells were also transfected with plasmids harboring human MIPS. (a) Western blot analysis was performed to confirm successful knockout of ISYNA1 and proper expression of ISYNA1 from the rescue plasmid. (b) Translation alignment of WT and Cas9-edited (KO) ISYNA1 alleles following Sanger sequencing. Note the presence of frameshift mutations and a premature stop codon in both ISYNA1-KO alleles.

### 3.2. Inositol deprivation induces cell death

To characterize the requirement for inositol, ISYNA1-KO cells were cultured in inositol-free DMEM supplemented with 10% dialyzed fetal bovine serum (FBS). Surprisingly, the ISYNA1-KO cells had no observable phenotypic difference from WT cells under these conditions (data not shown). This suggested that the serum, although dialyzed, contained enough inositol to support cell growth. Analysis of inositol content confirmed the presence of inositol (~25 μM) in commercial dialyzed FBS [39]. Therefore, the serum was further dialyzed using 3.5 kDa MWCO dialysis tubing. The absence of inositol in the dialyzed serum was confirmed using HPLC-MS [39]. Furthermore, a novel inositol bioassay confirmed that the lab-dialyzed serum, unlike the commercially available product, does not support the growth of the well-characterized inositol auxotrophic yeast strain *ino1*Δ [39].

To establish whether inositol-less death occurs in human cells, cell viability was assayed using inositol-free DMEM supplemented with 10% lab-dialyzed FBS. ISYNA1-KO cells showed decreased viability after four days of inositol depletion, which resulted in the complete death of cultures by day six (Fig. 2a). Interestingly, in the absence of exogenous inositol, WT cells showed reduced proliferation after four days of growth, demonstrating that inositol uptake is essential for maintaining the optimal proliferation of these cells (Fig. 2a). To characterize the specific effect of MIPS on cell survival, an ISYNA1 expression plasmid was constructed by cloning ISYNA1 cDNA [28] into a constitutive expression plasmid downstream of the human cytomegalovirus (CMV) promoter. Western blot analysis confirmed the restoration of MIPS protein in KO cells transfected with the expression plasmid (Fig. 1a), which rescued the inositol-less death phenotype observed in inositol-free medium (Fig. 2b).

**Figure 2.**
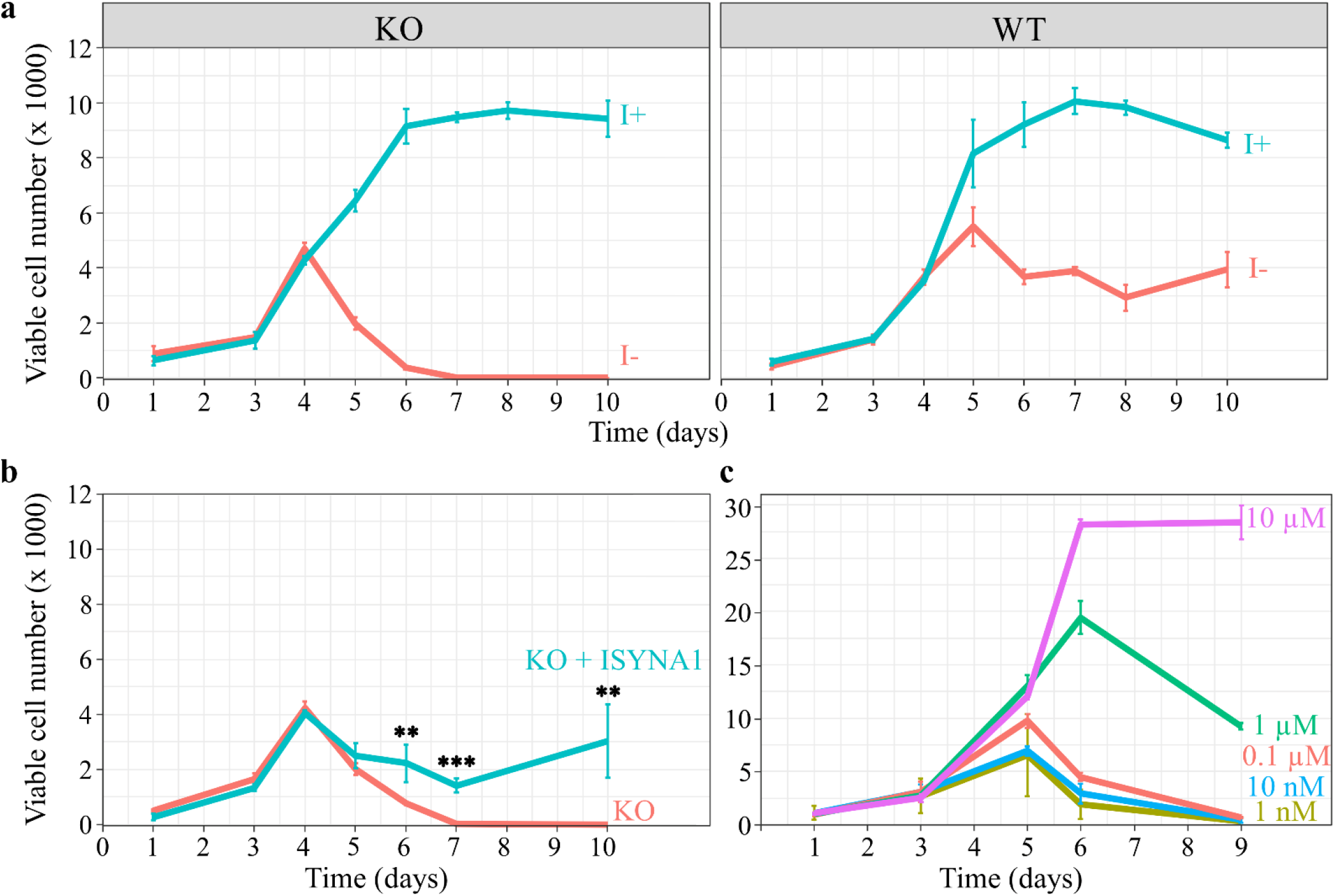
Inositol depletion leads to cell death. HEK293T WT and KO cells were cultured in 96-well plates, and viable cell numbers were estimated by the viability assay described in ‘Materials and methods.’ (a) Inositol (1 mM) promotes the growth of WT and KO cells, while inositol deprivation decreases the viability of KO cells after four days, eventually leading to total cell death by day 6. Water was used as a vehicle. (b) Overexpression of ISYNA1 rescues viability of ISYNA1-KO cells when cultured in inositol-free medium. T-tests were performed to infer significance (**, *p* ≤ 0.01; ***, *p* ≤ 0.001). (c) A range of inositol concentrations was used to determine the minimum concentration requirements to sustain cell viability. The data shown are the average of at least three replicates ± SD, n ≥3. The seeding densities for (a) and (b) are ~700 cells/well, and ~1000 cells/well for (c).

To determine the minimum inositol concentration that permits cell proliferation, ISYNA1-KO cells were cultured in media supplemented with a series of ten-fold serial dilutions of *myo*-inositol, the most abundant of nine inositol isomers [27]. The minimum inositol concentration that permitted growth was determined to be 10 μM (Fig. 2c).

Aside from *myo*-inositol, two of the most studied inositol isomers are *scyllo*-inositol and D-*chiro*-inositol, which are associated with human diseases such as Alzheimer’s disease, diabetes, and polycystic ovary syndrome [27]. To test whether these additional isomers could support cell growth in the absence of *myo*-inositol, ISYNA1-KO cells were cultured in inositol-free medium supplemented with 500 μM *scyllo*-inositol or D-*chiro*-inositol. Interestingly, neither of these isomers could compensate for the absence of *myo*-inositol, as they did not support the growth of ISYNA1-KO cells (Fig. 3a). Similarly, supplementation of *scyllo*-inositol or D-*chiro*-inositol (100 μM) did not support the growth of yeast *ino1*Δ when cultured in inositol-free medium (Fig. 3b), demonstrating that the cellular requirement for inositol is specific to the *myo*isomer.

**Figure 3.**
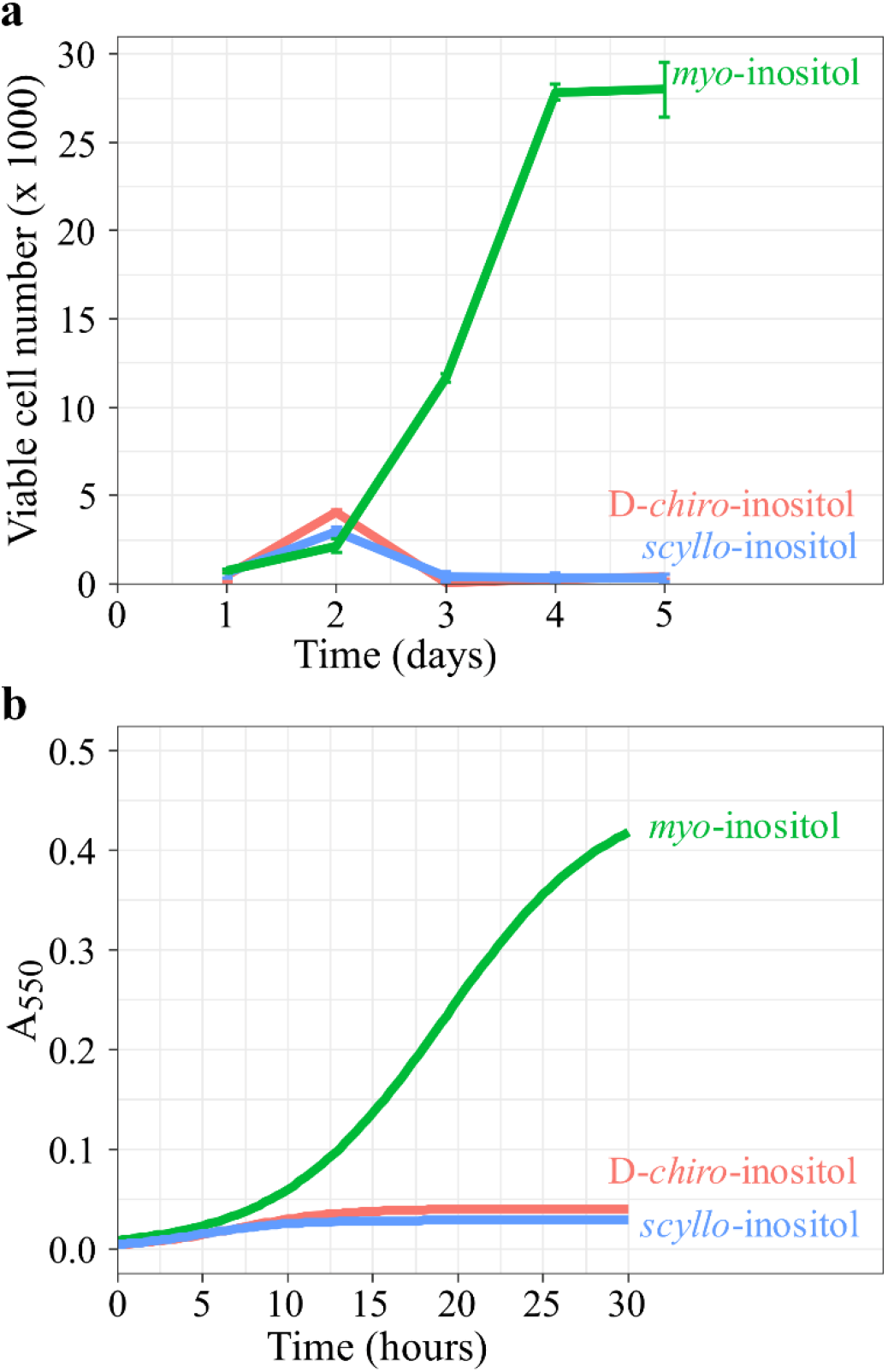
D-*chiro*-inositol and *scyllo*-inositol do not support viability. (a) HEK293T ISYNA1-KO cells were cultured in 96-well plates (~1000 cells/well) in inositol-free medium supplemented with either 10 uM inositol, 500 uM D-*chiro*-inositol, or 500 uM *scyllo*-inositol. Viable cell numbers were estimated using the viability assay described in ‘Materials and methods’. (b) *ino1*Δ cells were inoculated at A_550_ 0.05 in 180 μL of liquid SCD medium in a 96-well plate supplemented with either 10 uM inositol, 100 uM D-*chiro*-inositol, or 100 uM *scyllo*-inositol. Cells were incubated at 30°C and A_550_ was determined at 20 min intervals until stationary phase using a SpectraMax i3x plate reader (Molecular Devices, San Jose, CA), running an automated growth curve protocol. The data shown are the average of at least three replicates ± SD, n ≥3.

The finding that inositol-less death is conserved from yeast to human cells suggested that other aspects of inositol regulation may be highly conserved. Studies in yeast have demonstrated that *INO1*, the yeast homolog of ISYNA1, is the most responsive gene to external inositol, as its expression is upregulated substantially in the absence of inositol [18]. Therefore, to determine whether ISYNA1 gene expression is regulated by inositol, Western blot analysis of WT HEK293T MIPS was performed in the presence or absence of external inositol. Surprisingly, this showed that inositol does not regulate MIPS protein levels in human cells (Fig. 4a). This is in striking contrast to yeast MIPS for which expression is almost completely abolished in the presence of inositol (Fig. 4b).

**Figure 4.**
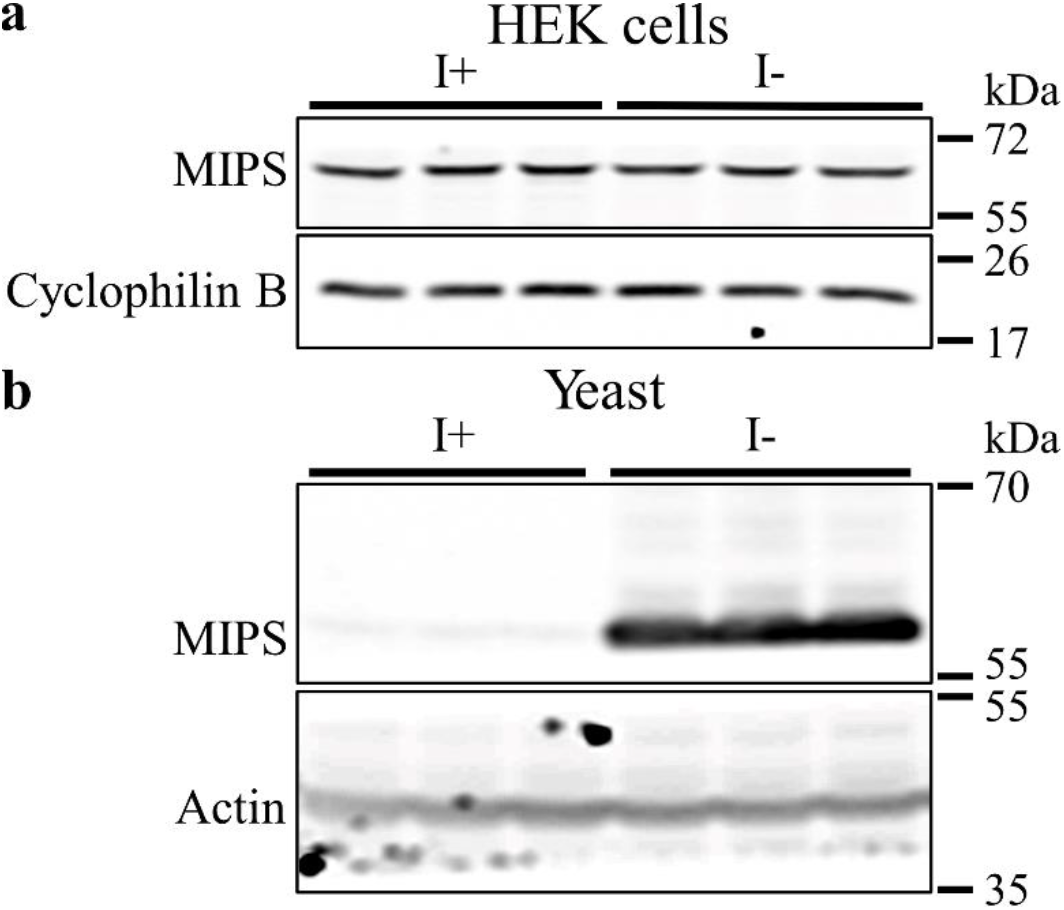
Exogenous inositol does not regulate MIPS protein levels in WT HEK293T cells. (a) WT cells were cultured for four days in an inositol-free medium supplemented with 1 mM inositol. Western blot analysis was performed using an antibody against human MIPS, and cyclophilin B was used as a loading control. (b) WT BY4741 yeast cultures were grown in triplicate to mid-log phase in SCD I+ medium. Cultures were then split in two, washed once with water, then resuspended in SCD I+ (75 μM) or SCD I- and incubated for 4 hrs before collecting cell extracts. Western blot analysis was performed using a custom antibody against yeast MIPS, and actin was used as a loading control.

### 3.3. Inositol deprivation induces significant changes in phospholipid metabolism

Inositol is a primary component of the phospholipid PI, and inositol depletion has been shown to regulate phospholipid levels in yeast and Chinese hamster ovary (CHO) cells [40–42]. To determine if inositol regulates phospholipid metabolism in human cells, phospholipid profile analysis of ISYNA1-KO cells was performed after four days of inositol depletion using HPLC-MS (Table S2). As expected, the analysis demonstrated that inositol deprivation resulted in more than a three-fold decrease in PI levels (Figs. 5-6). Phosphatidic acid (PA) levels were also downregulated, and phosphatidylethanolamine (PE) levels were upregulated following inositol depletion, whereas there was no significant effect on total phosphatidylcholine (PC) or phosphatidylserine (PS) (Figs. 5-6). Remarkably, a robust (~10-fold) increase in PG levels was observed, as well as significant upregulation of phospholipid species synthesized from PG, including CL, monolysocardiolipin (MLCL), dilysocardiolipin (DLCL), and bis(monoacylglycerol)phosphate (BMP) (Fig. 5 and Fig. 7). This analysis indicates that inositol depletion strongly affects the lipidome of mitochondria, where PG and CL are synthesized [43].

**Figure 5.**
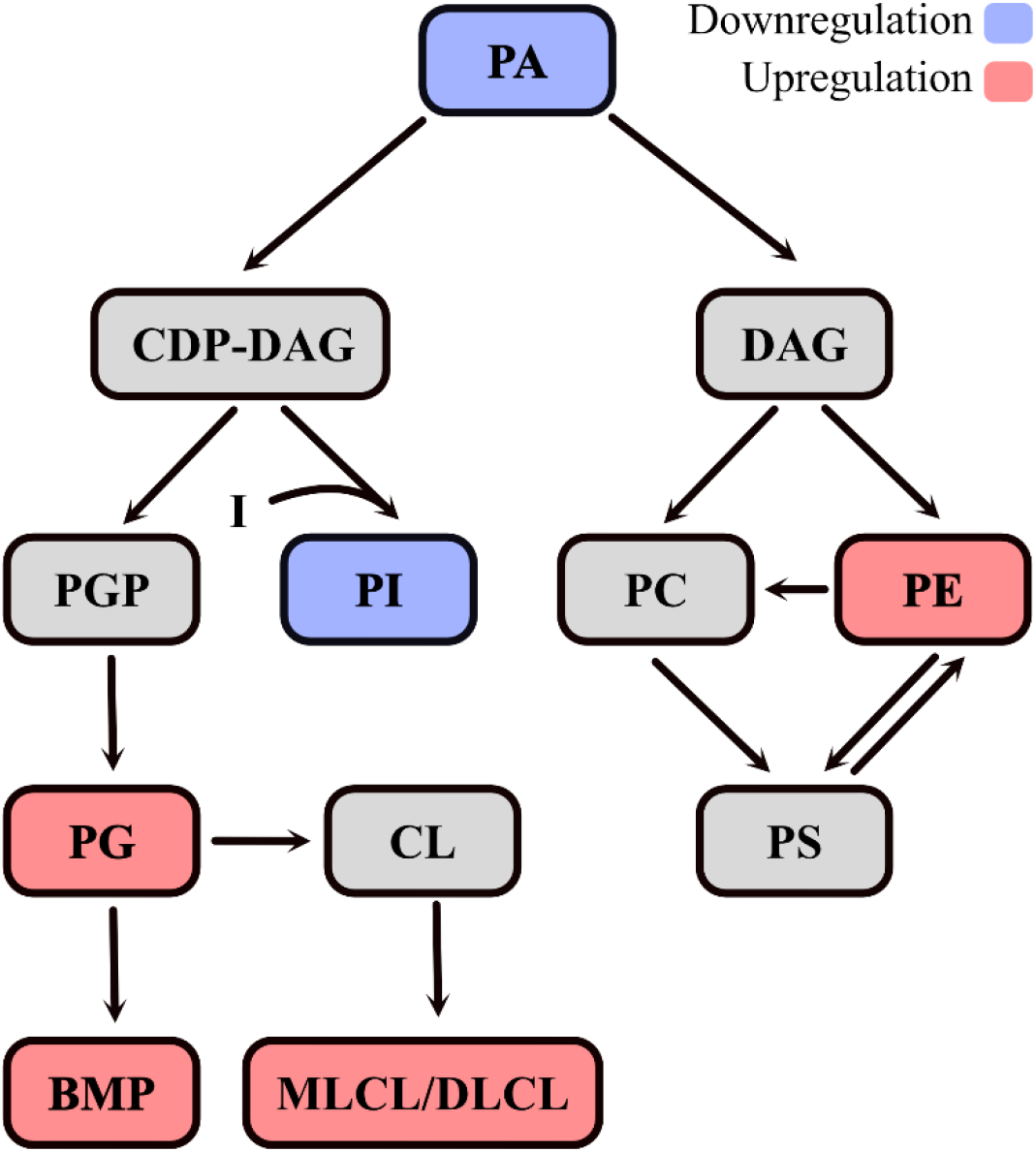
The mammalian phospholipid pathway depicting the effects of inositol depletion. The diagram represents a simplified depiction of the phospholipid metabolic pathway and the differentially regulated phospholipids resulting from inositol deprivation. Red color indicates upregulation while blue color indicates downregulation. PA is converted into CDP-DAG and DAG which are the precursors of phospholipids. PI is generated in the ER from CDP-DAG and inositol, while PGP is synthesized in mitochondria from CDP-DAG and glycerol-3-phosphate. PGP is further dephosphorylated to PG, which is converted to CL. BMP is generated from PG in a complex multi-step reaction. PC and PE are generated by combining DAG with CDP-choline or CDP-ethanolamine, respectively. PS can be synthesized from PC or PE. Additional interconversion steps between PC, PE, and PS are indicated.

**Figure 6.**
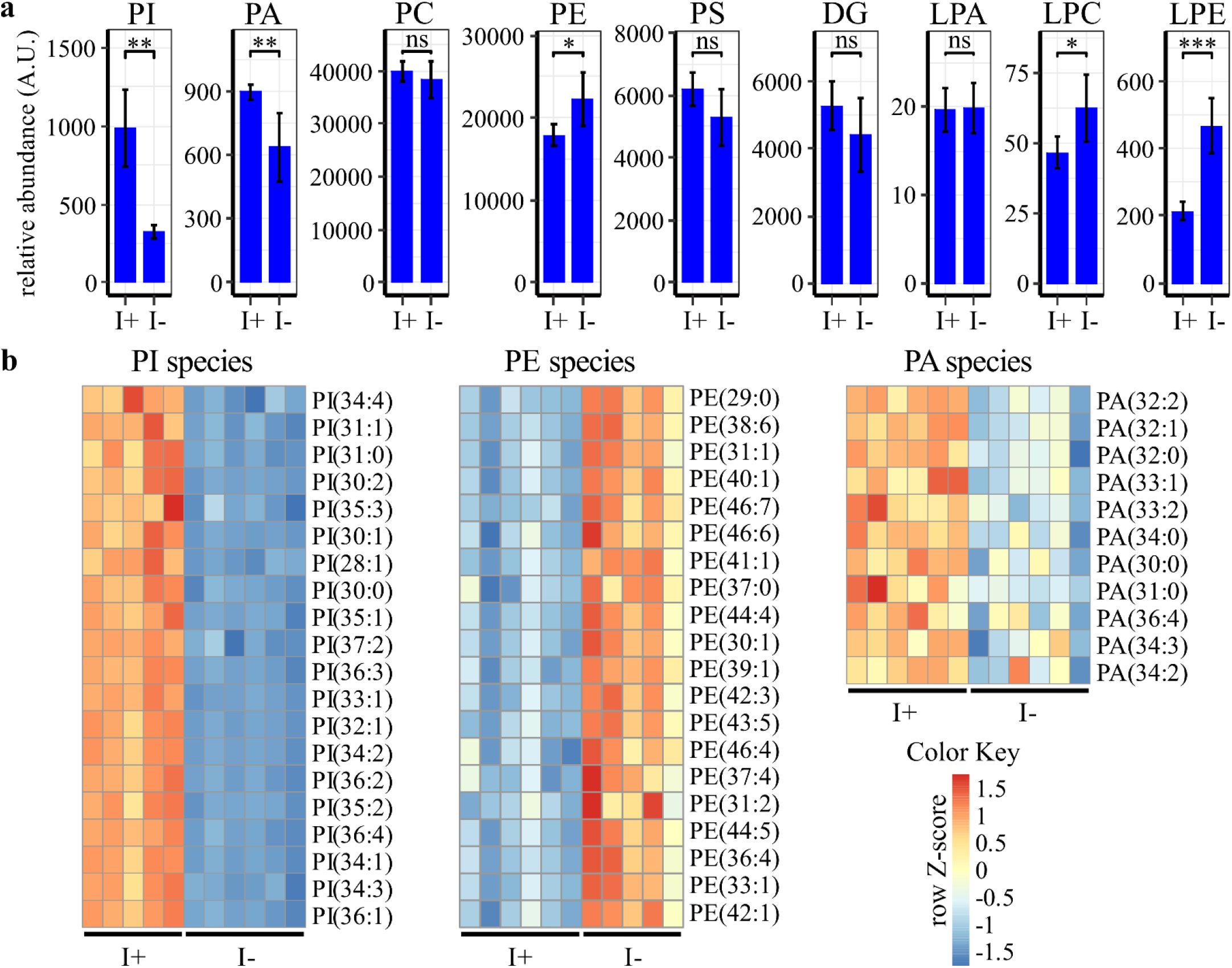
Inositol depletion downregulates PI and PA levels and upregulates PE levels. ISYNA1-KO cells were cultured for four days in inositol-free medium. Cells were washed with PBS, lysed, then phospholipids extracted and analyzed using HPLC-MS. Data were normalized to the protein content of each sample. (a) Total phospholipid levels are shown as the average ± SD, n ≥ 5. A Student’s t-test was utilized to determine significance (*, p ≤ 0.05; **, p ≤ 0.01; ***, p < 0.001). (b) Heatmaps of significantly regulated PI, PE, and PA species were generated using row Z-scores of relative abundance units for each species. Species are ranked based on significance (p ≤ 0.05), while only the top 20 are included for PI and PE. LPC = lysophosphatidylcholine, LPE = Lysophosphatidylethanolamine; LPA = Lysophosphatic acid.

**Figure 7.**
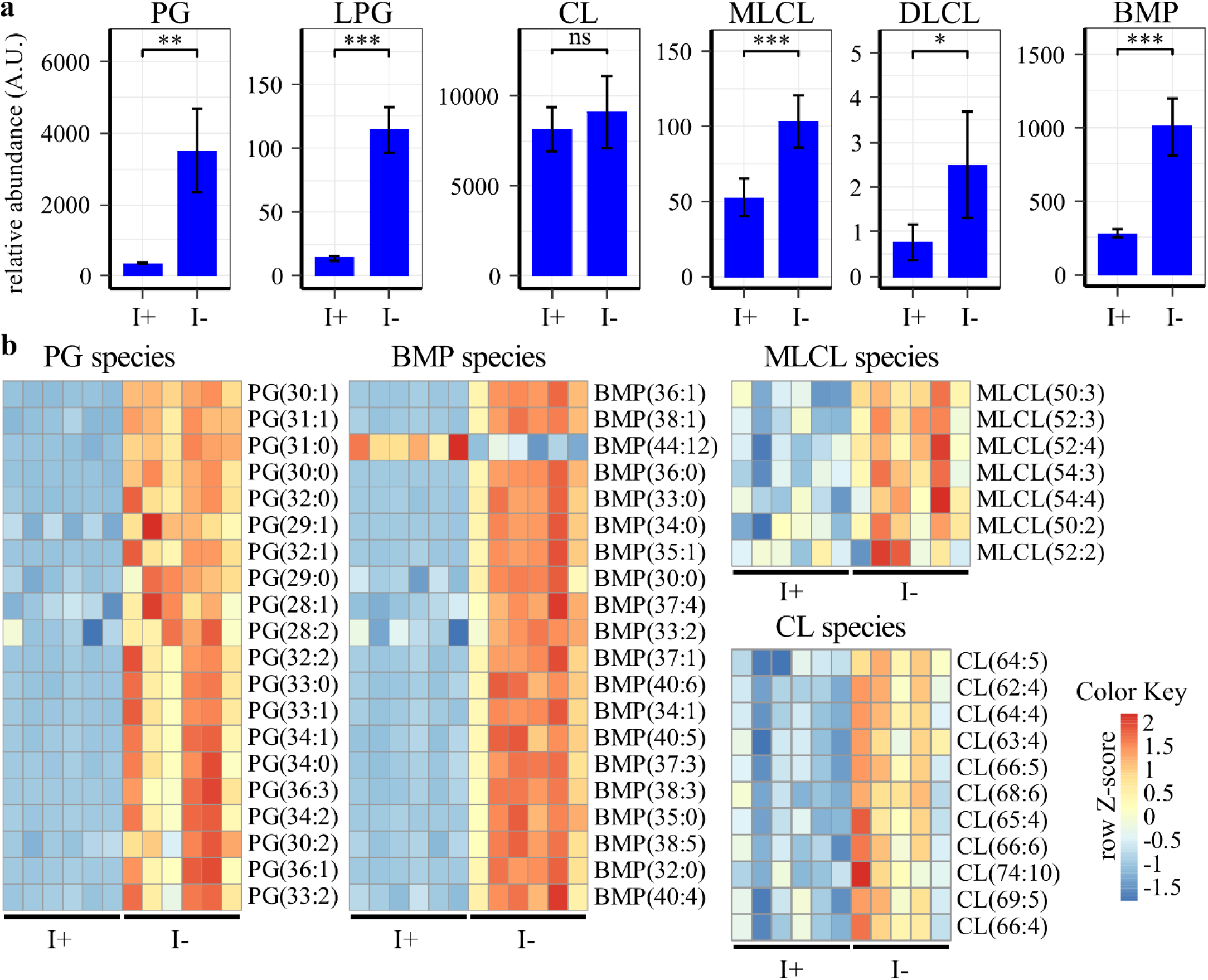
Inositol depletion upregulates PG/CL-derived lipids. ISYNA1-KO cells were cultured for four days in inositol-free medium. Cells were washed with PBS, lysed, then phospholipids extracted and analyzed using HPLC-MS. Data were normalized to the protein content of each sample. (a) Total phospholipid levels are shown as the average ± SD, n ≥ 5. A Student’s t-test was utilized to determine significance (*, p ≤ 0.05; **, p ≤ 0.01; ***, p < 0.001). (b) Heatmaps of significantly regulated PG, BMP, CL, and MLCL species were generated using row Z-scores of relative abundance units for each species. Species are ranked based on significance (*p* ≤ 0.05), while only the top 20 are included for PG and BMP.

### 3.4. Inositol deprivation induces profound changes in gene transcription, including ERK signaling and stress response genes

The changes in lipid metabolism described above suggested that inositol deprivation may induce widespread changes in gene transcription. To further characterize the pathways regulated by inositol deprivation, RNA-Seq analysis of ISYNA1-KO cells was performed after four days of inositol depletion (Table S3). Inositol deprivation induced profound changes in gene transcription, significantly upregulating a total of 191 genes while downregulating 108 genes (Fig. 8). Gene enrichment analysis of the differentially expressed genes identified mitogen-activated protein kinase (MAPK) signaling, amino acid metabolism, PPAR signaling, steroid biosynthesis, and endoplasmic reticulum protein processing among the most highly regulated pathways (Fig. 9). Surprisingly, there were only minor effects on phospholipid metabolism genes, with only four out of ~100 exhibiting differential expression (Fig. S1 and Table S5).

**Figure 8.**
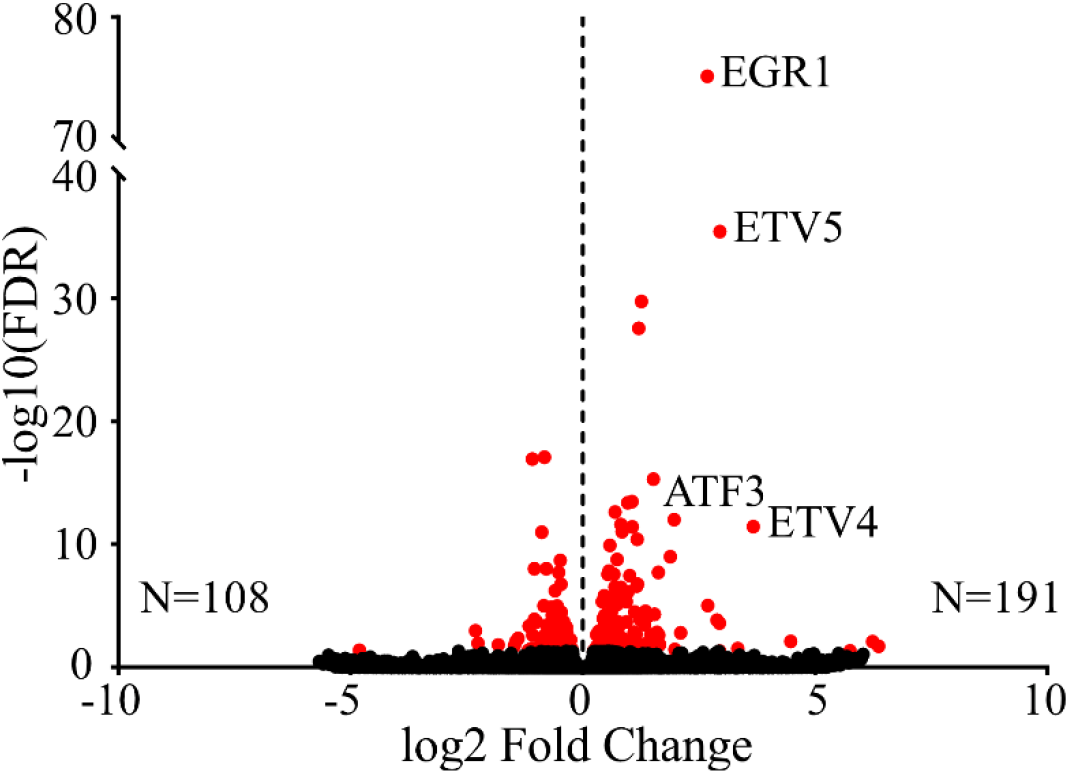
Inositol depletion induces broad changes in gene transcription. ISYNA1-KO cells were cultured for four days in inositol-free medium. The volcano plot depicts log2 fold change and -log10 (FDR) of differentially expressed genes. Significantly upregulated and downregulated transcriptional changes are depicted by red dots with the total number of genes in each category indicated (FDR ≤ 0.05).

**Figure 9.**
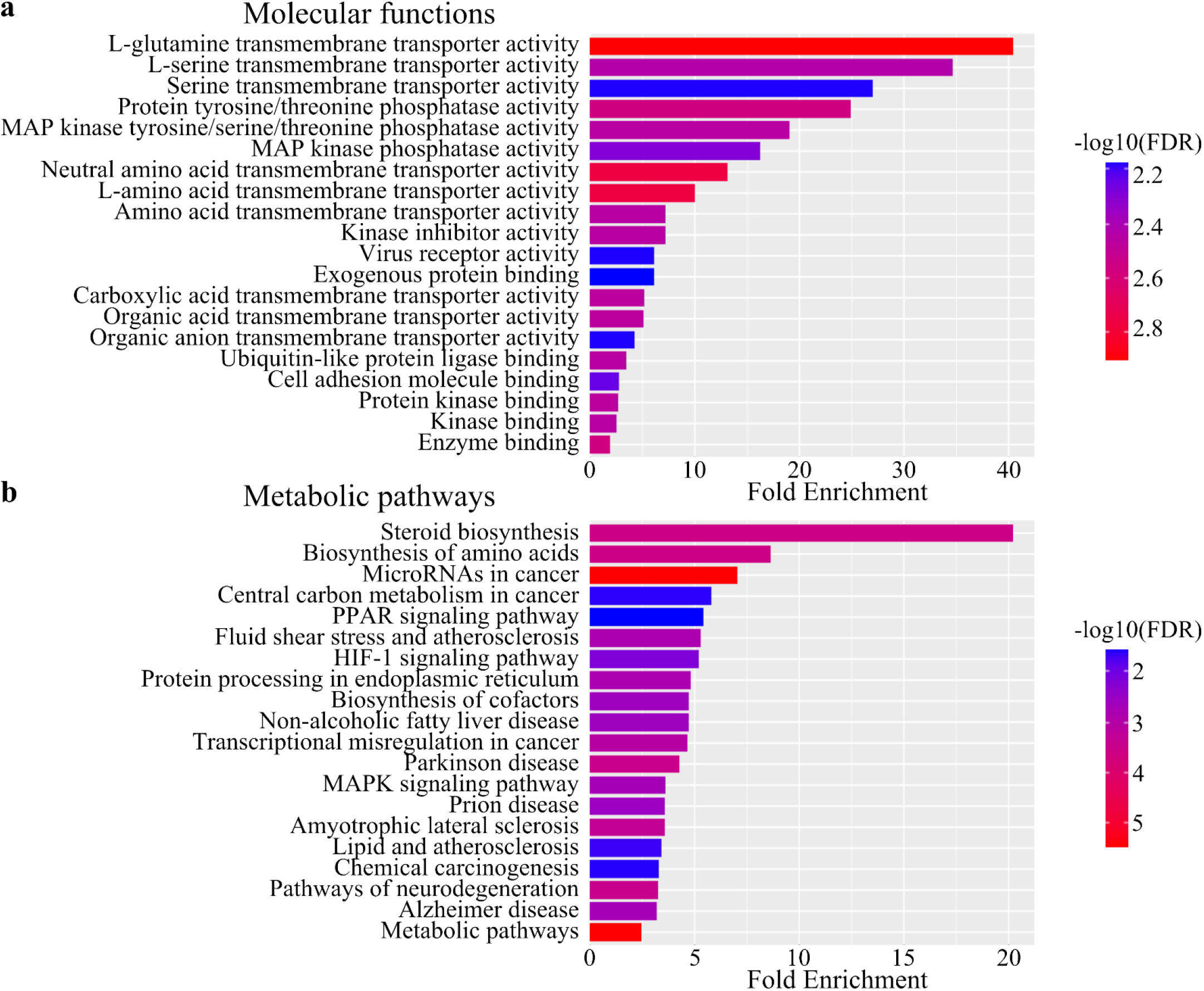
Inositol depletion induces significant transcriptional changes in signaling and metabolic pathways. ISYNA1-KO cells were cultured for four days in inositol-free medium. Gene enrichment analysis of differentially expressed genes was performed using ShinyGO (v0.741). The enrichment was performed for (a) GO molecular functions and (b) Kyoto Encyclopedia of Genes and Genomes (KEGG) pathways. The results are ranked based on fold enrichment and FDR.

The gene enrichment analysis identified MAPK signaling among the most highly upregulated pathways (Fig. 9). Further inspection of the data revealed that a cluster of genes regulated by extracellular signal-regulated kinase 1 (ERK1) and ERK2 signaling are particularly elevated in expression (Fig. 10a, and table S1). RT-PCR analysis of two of these genes, EGR1 and ETV4, confirmed this observation (Fig. S2). Western blot analysis was performed to determine whether ERK signaling is activated under inositol depletion using antibodies against phosphorylated ERK, a marker for activation of the corresponding kinase cascade. The results show that inositol depletion increased the phosphorylation of ERK1 and ERK2 by four-fold (Fig. 10b), suggesting that the ERK signaling cascade responds to inositol depletion.

**Figure 10.**
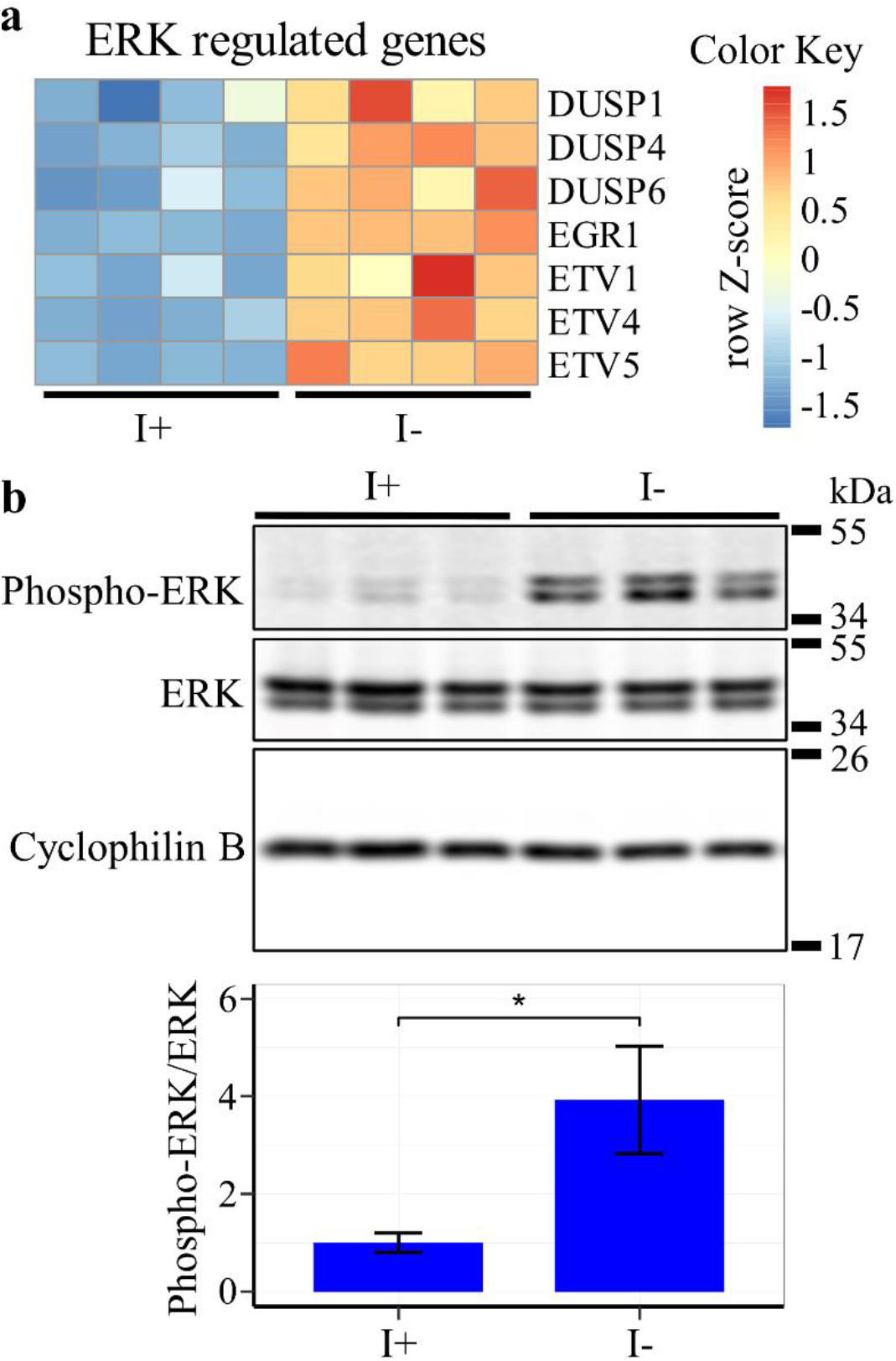
Inositol depletion upregulates ERK. ISYNA1-KO cells were cultured for four days in inositol-free medium. (a) A heatmap of significantly affected ERK-regulated genes was generated using row Z-scores of normalized counts for four replicates for each condition (FDR ≤ 0.05). (b) Western blot analysis was performed using an antibody against Phospho-ERK and total ERK protein, and cyclophilin B was used as a loading control. Band intensity was quantified using ImageJ software. Data shown are average fold change of Phospho-ERK/ERK ± SD, n = 3 (*, p ≤ 0.05).

In addition to ERK, the enrichment analysis also indicated that genes involved in cellular stress response signaling downstream of activating transcription factor 4 (ATF4) and various amino acid transporters are among the most upregulated in response to inositol deprivation (Fig. 11a and table S1). Western blot analysis confirmed that ATF4 protein levels are elevated under these conditions (Fig. 11b), indicating activation of the pathway. The data above suggest that both ERK and ATF4 signaling cascades respond to the metabolic and nutrient stress induced by inositol deprivation. Cells adapt to metabolic stress and nutrient limitation by inducing autophagy [44]. One of the primary hallmarks of autophagy is the cleavage of microtubule associated protein 1 light chain 3 beta-I (LC3B-I) to produce LC3B-II during autophagosome formation and maturation [45]. Western blot analysis indicated an increase in the LC3B-II to LC3B-I ratio following inositol deprivation, suggesting induction of autophagy in this context (Fig. 12) [46]. In summary, inositol depletion causes robust activation of the ERK and ATF4 signaling pathways and upregulates autophagy, suggesting that these outcomes may play a role in the observed inositol-less death.

**Figure 11.**
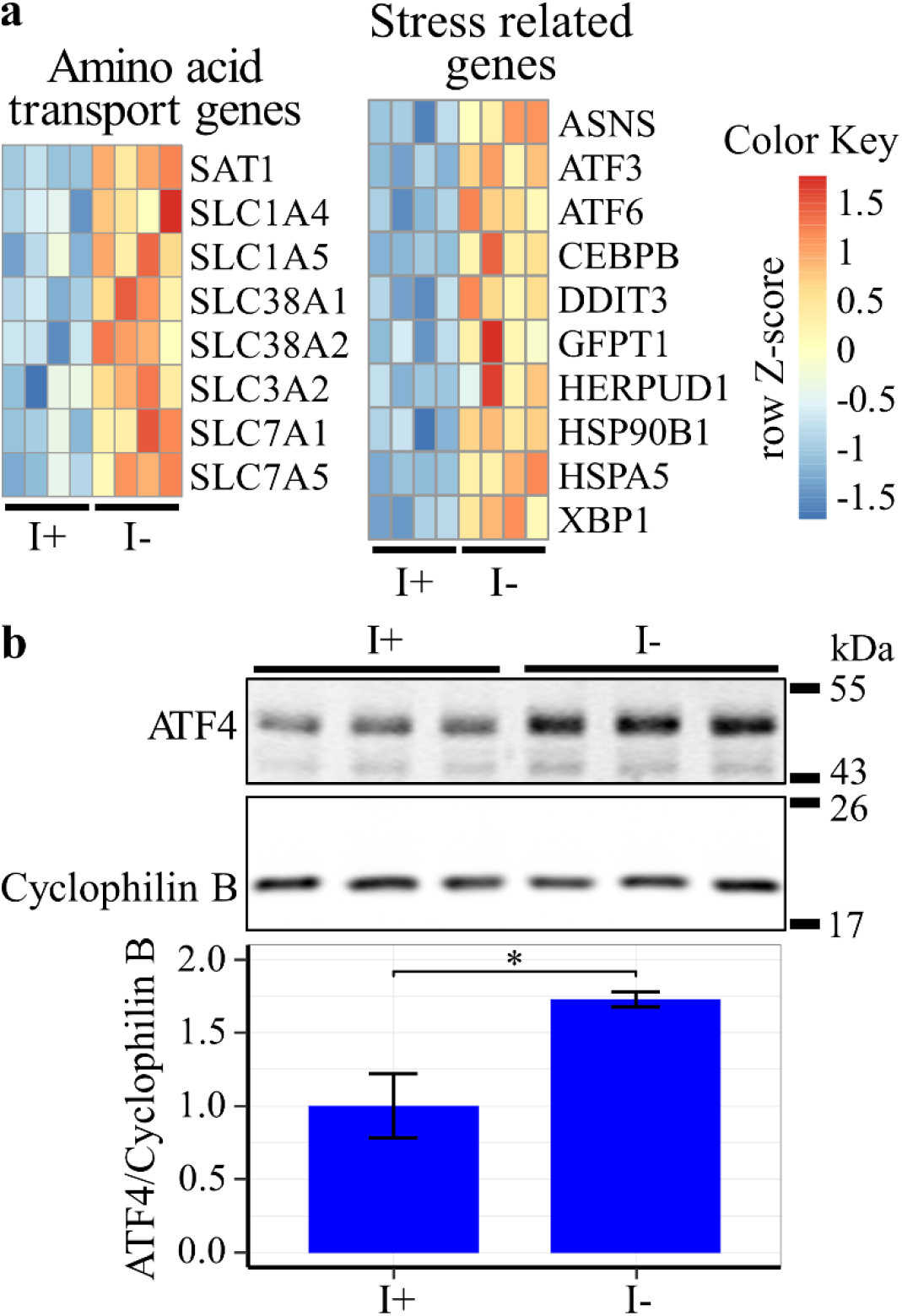
Inositol depletion upregulates ATF4. ISYNA1-KO cells were cultured for four days in inositol-free medium. (a) Heatmaps of significantly regulated amino acid transport genes and stress response genes were generated using row Z-scores of normalized counts for four replicates in each condition (FDR ≤ 0.05). (b) Western blot analysis was performed using an antibody against ATF4, and cyclophilin B was used as a loading control. Band intensity was quantified using ImageJ software. Data shown are average fold change of ATF4/Cyclophilin B ± SD, n = 3 (*, p ≤ 0.05).

**Figure 12.**
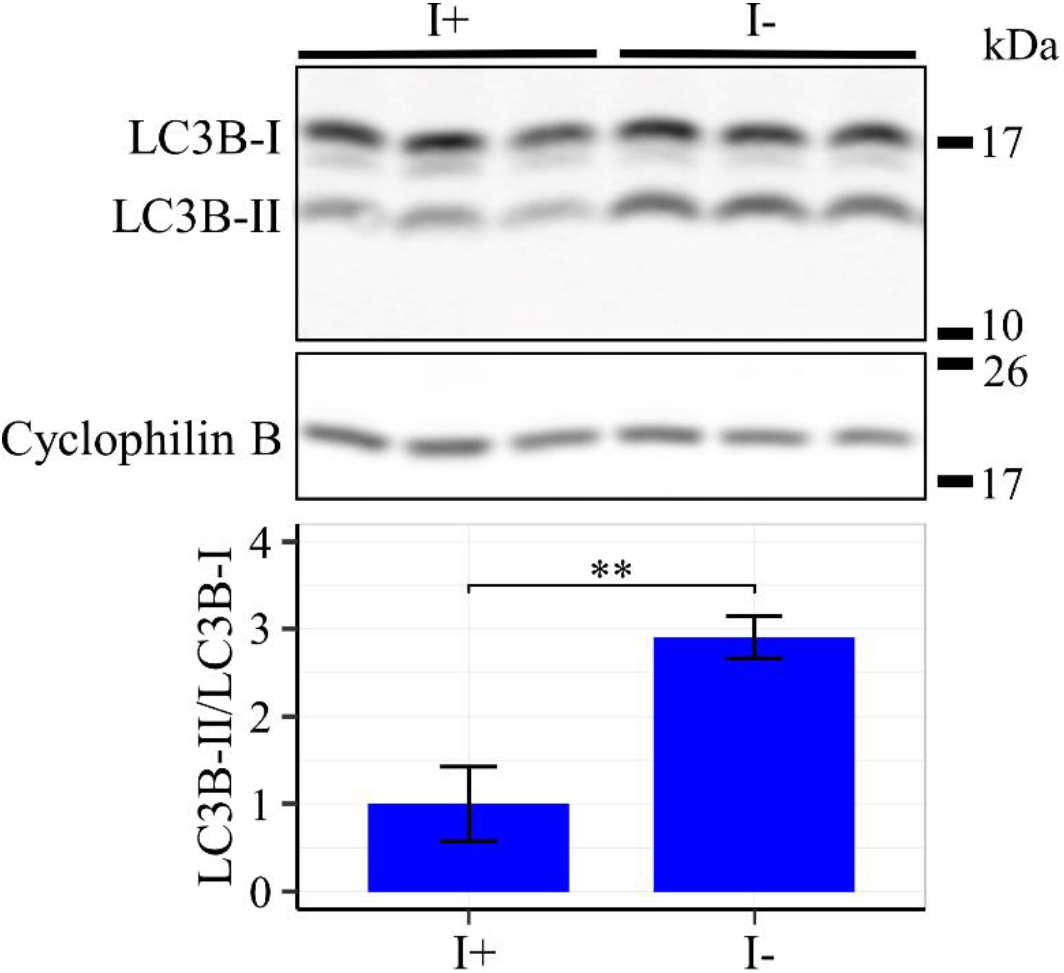
Inositol depletion upregulates LC3B-II/LC3B-I ratio. ISYNA1-KO cells were cultured for four days in inositol-free medium. Western blot analysis was performed using an antibody against LC3B protein, and cyclophilin B was used as a loading control. Band intensity was quantified using ImageJ software. Data shown are average fold change of LC3B-II/LC3B-I ± SD, n = 3 (**, p ≤ 0.01).

## 4. DISCUSSION

Inositol and inositol-containing compounds play an essential role in cell signaling and metabolism [2, 9]. Despite its importance, very little is known about inositol regulation in human cells [3]. This study provides the first in-depth characterization of the effects of inositol depletion on cell viability, phospholipid levels, and global gene regulation in a human cell line. Our findings demonstrate that inositol regulates a myriad of diverse cellular pathways but has a particularly robust influence on mitochondrial phospholipid metabolism and cellular stress response signaling.

It was previously demonstrated that either supplementation or *de novo* synthesis of inositol is sufficient to support cell mass accumulation of a colon cancer cell line [12]. The current study further illustrates that inositol is an essential metabolite for cell proliferation and viability. In the absence of supplemental inositol, ISYNA1-KO cells lose viability after four days (Fig. 2a). The observed loss in viability for this cell line can be directly attributed to the loss of MIPS function, as ectopic MIPS expression rescued the inositol-less death phenotype (Fig. 2b). The requirement for *myo*-inositol is specific, as D-*chiro-* and *scyllo*-inositol isomers could not rescue viability (Fig. 3). When inositol cannot be obtained from extracellular sources, WT human cells are able to rely on *de novo* inositol synthesis to remain viable. However, our data indicate that even WT cells require environmentally derived inositol for optimal cell growth, as WT cell proliferation was inhibited after four days of growth in inositol-free medium (Fig. 2a). Though commonplace in mammalian cell culture, FBS is often overlooked as a source of experimental variability. The finding that commercially available dialyzed FBS contains sufficient inositol to support the viability of ISYNA1-KO cells highlights the importance of controlling for batch-to-batch variation in FBS composition, which could result in widespread changes in gene regulation based on our findings.

Yeast cells tightly regulate MIPS expression in response to external inositol (Fig. 4b) [3]. This is in striking contrast to WT HEK293T cells, where external inositol had no effect on expression of MIPS protein (Fig. 4a) or mRNA levels (data not shown). This finding is corroborated by a previous study in a neuronal cell line that showed ISYNA1 mRNA and MIPS protein levels were unaffected by external inositol [47]. These findings indicate that ISYNA1 is regulated by a different mechanism compared to its *INO1* homolog in yeast. A candidate mechanism for MIPS regulation in mammalian cells is transcriptional repression by inositol hexakisphosphate kinase 1 (IP6K1), as was demonstrated in mouse embryonic fibroblasts [49]. In the corresponding study, it was shown that IP6K1 translocates to the nucleus in a PA-dependent manner, binds to the ISYNA1 promotor, and downregulates ISYNA1 transcription by inducing hypermethylation [49]. Intriguingly, negative regulation of ISYNA1 transcription by hypermethylation has also been demonstrated in human acute myeloid leukemia (AML) cells [21]. However, the current study indicates that depletion of inositol itself leads to a decrease in PA levels without affecting MIPS protein abundance, suggesting that additional details of this mechanism have yet to be elucidated.

Inositol depletion has a profound effect on phospholipid metabolism. Lipidomic analysis demonstrated that inositol depletion decreases PI levels more than three-fold (Fig 6). This suggests that signaling mediated by phosphoinositides, which are synthesized from PI, may also be reduced due to a lack of biosynthetic substrate [9]. In agreement with this, studies have shown that inositol depletion decreases phosphoinositide levels in yeast and CHO cells [17]. This is also consistent with a previous *in vivo* study that demonstrated a significant decrease in intestinal microsomal PI levels when gerbils were fed an inositol-deficient diet [50]. The lipidomic analysis also indicated a decrease in PA levels (Fig. 6). Although a decrease in PI levels would be expected to lead to a concomitant increase in PA, as was demonstrated in yeast [51], the observed decrease in PA may instead be the result of increased PG and PE synthesis. PG synthesis consumes PA through CDP-DAG [52], whereas PE synthesis consumes PA through DAG (Fig. 5). Therefore, further analysis of these phospholipid intermediates may clarify the exact fate of PA following inositol depletion. Notably, RNA-Seq analysis indicated that transcription of most phospholipid biosynthetic genes is not significantly altered by inositol depletion (Fig. S1 & Table S5), suggesting that the observed changes in phospholipid levels depend more on differences in enzymatic activity or substrate availability.

Inositol depletion exerts differential effects on total PE and PC levels. The lipidomic data indicated upregulation of total PE levels whereas there was no significant change in total PC or PS levels (Fig 6). Interestingly, an increase in PE without a corresponding change in PC levels was also observed in the intestinal microsome of gerbils fed an inositol-deficient diet [50, 53]. One means of *de novo* synthesis of PC and PE in eukaryotes can occur *via* the Kennedy pathway [54]. The rate-limiting enzymes in the respective biosynthetic component pathways are CTP:phosphocholine cytidylyltransferase (CCT) encoded by the PCYT1 gene in the PC pathway and CTP:phosphoethanolamine cytidylyltransferase (ET) encoded by the PCYT2 gene in the PE pathway [54]. Our RNA-Seq data indicated a decrease in PCYT2 transcription while PCYT1 was unchanged (Fig. S1, and table S5). Therefore, our findings cannot be explained by transcriptional regulation of these enzymes. An alternative explanation for this is that enzymatic activity of ET is increased following inositol deprivation while CCT activity is unaffected in these conditions. PE can also be synthesized by decarboxylation of PS and subsequently converted to PC by methylation [55]. Therefore, future studies examining the relative activities of the interconverting enzymes in these highly complex pathways are needed to fully elucidate the mechanism underlying the current finding.

Strikingly, inositol depletion also resulted in a ten-fold increase in PG levels and a two-fold or greater increase in phospholipid species downstream of PG, including CL, MLCL, DLCL, and BMP (Fig. 7). The observed increase in PG following inositol deprivation is in agreement with a previous study in CHO cells [42]. Interestingly, in yeast, inositol inhibits PGS1, a key enzyme in PG synthesis [56]. This suggests that a similar mechanism of PGS1 inhibition could exist in human cells. PG is synthesized in mitochondria where it serves as the precursor for CL and BMP [57]. MLCL and DLCL, intermediates in the CL remodeling pathway, play a significant role in maintaining the structure and function of mitochondrial membranes, while BMP is essential for cholesterol metabolism and trafficking in lysosomes [58–60]. Therefore, these findings suggest that inositol regulation may be linked to mitochondrial integrity and lysosome function through alterations in phospholipid metabolism.

RNA-Seq analysis showed that inositol depletion induces widespread changes in gene expression related to ERK signaling, amino acid metabolism, stress signal transduction, and steroid biosynthesis (Fig. 9 and table S3). Pathways regulated by ERK represent the highest upregulated cluster of genes, and activation of ERK was further demonstrated by increased phosphorylation of the protein (Fig. 10). ERK is part of the MAPK signaling cascade and plays a significant role in cell proliferation, metabolism, migration, differentiation, and survival [61, 62]. Regulation of ERK by inositol may have therapeutic implications. For example, a previous report identified inositol as a potential treatment for lung cancer through ERK-mediated inhibition of bronchial lesion proliferation [63].

ERK has also been shown to mediate the cellular response to metabolic and nutrient stress by increasing the expression of ATF4 [64]. ATF4 is an effector of the integrated stress response network, comprised of several stress response pathways that respond to amino acid starvation, heme deprivation, and ER stress [65]. Upon activation, ATF4 restores homeostasis by activating genes related to the unfolded protein response, oxidative stress, and amino acid transport [65, 66]. Gene enrichment analysis indicated upregulation of genes regulated by ATF4, including amino acid transporters and stress response genes in response to inositol deprivation (Fig. 11a and Table S1). Western blot confirmed increased expression of ATF4 protein levels (Fig. 11b). This suggests that cells identify inositol deprivation as a form of nutrient stress and respond by activating the integrated stress response. One way that cells counteract nutrient limitation is by inducing autophagy [44], and this was demonstrated to occur in response to inositol deprivation (Fig. 12). Collectively, these findings indicate that inositol is a major regulator of ERK signaling and the integrated stress response.

In conclusion, this study provides the first extensive characterization of the cellular consequences of inositol deprivation in human cells. Our results demonstrate that inositol is essential for maintaining cell viability and that inositol deprivation is associated with dramatic changes to the phospholipidome and transcriptome. Furthermore, these findings highlight the importance of future studies aimed at elucidating the downstream effects of treatment with inositol-depleting drugs.

## The abbreviations used are

MIPS: *mvo*-inositol phosphate synthase
ISYNA1: inositol-3-phosphate synthase (mammalian)
*INO1*: inositol-3-phosphate synthase (yeast)
PI: phosphatidylinositol
PG: phosphatidylglycerol
PS: phosphatidylserine
PE: phosphatidylethanolamine
PC: phosphatidylcholine
CL: cardiolipin
MLCL: monolysocardiolipin
DLCL: dilysocardiolipin
BMP: bismonoacylglycerophosphate
DAG: diacylglycerol
CDP-DAG: cytidine diphosphate diacylglycerol
PGP: phosphoglycolate phosphatase
LPG: lysophosphatidylglycerol
LPE: lysophosphatidylethanolamine
LPC: lysophosphatidylcholine
CHO: Chinese hamster ovary cells
ERK: extracellular signal-regulated kinase
LC3B: microtubule associated protein 1 light chain 3 beta
FDR: false discovery rate
ER: endoplasmic reticulum
MAPK: mitogen-activated protein kinase
WT: wild-type
ATF4: activating transcription factor 4
LPC: lysophosphatidylcholine
LPE: Lysophosphatidylethanolamine
LPA: Lysophosphatic acid
FBS: fetal bovine serum
I3P: inositol-3-phosphate
CCT: CTP:phosphocholine cytidylyltransferase
ET: CTP:phosphoethanolamine cytidylyltransferase

## CRediT author statement

**Mahmoud Suliman:** Conceptualization, Methodology, Investigation, Writing - Original Draft, Visualization.

**Kendall C. Case:** Conceptualization, Methodology, Investigation, Writing - Review & Editing, Visualization.

**Michael W. Schmidtke:** Conceptualization, Methodology, Investigation, Resources, Writing - Review & Editing. **Pablo Lazcano:** Investigation, Writing - Review & Editing.

**Chisom J. Onu:** Investigation, Writing - Review & Editing.

**Miriam L. Greenberg**: Supervision, Funding acquisition, Conceptualization, Writing - Original Draft.

## Data statement

All data are contained within the article and supporting information.

## Acknowledgments

This work was supported by the National Institutes of Health grant number R01 GM125082 (to M.L.G) and a Rumble Fellowship from the Department of Biological Sciences, Wayne State University (to M.S.).

## Supplementary material for

**Figure S1.**
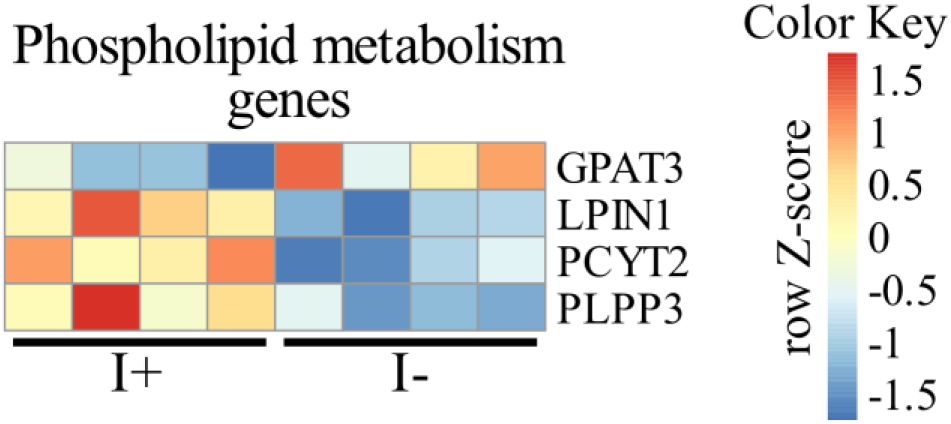
Phospholipid metabolism genes regulated by inositol depletion. ISYNA1-KO cells were cultured for four days in inositol-free medium. A heatmap of significantly regulated phospholipid metabolism genes was generated using row Z-scores of normalized counts for four replicates in each condition (FDR ≤ 0.05). Only four out of ~100 phospholipid metabolism genes were significantly altered by inositol deprivation.

**Figure S2.**
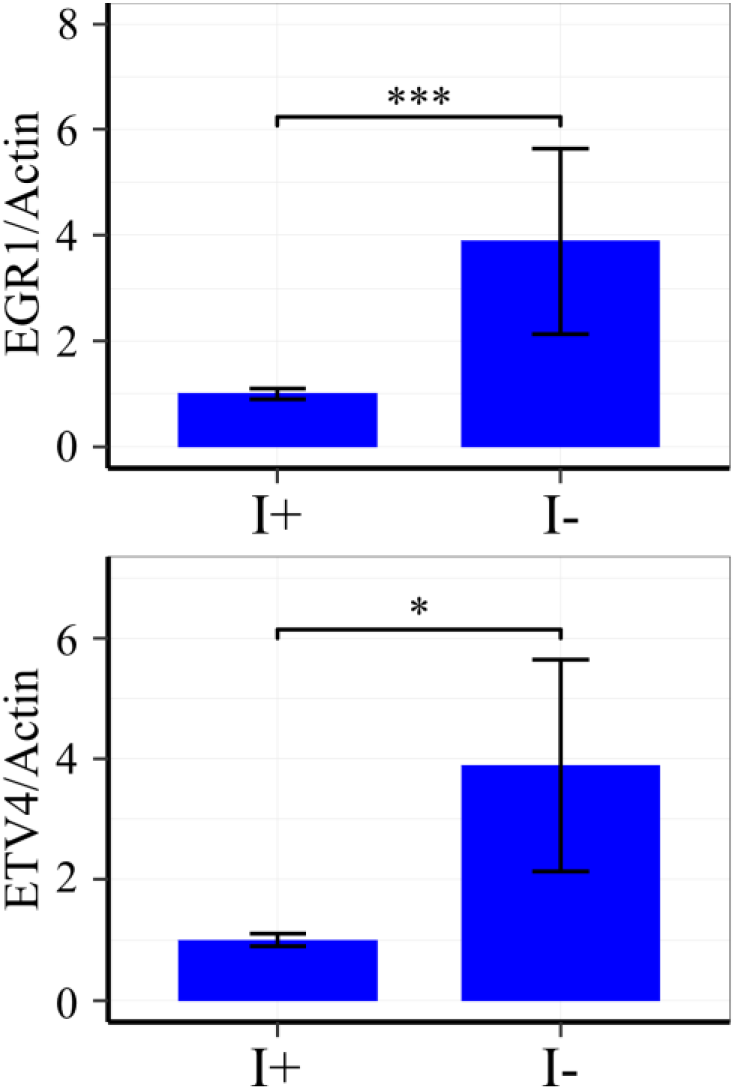
Inositol depletion upregulates EGR1 and ETV4. ISYNA1-KO cells were cultured for four days in inositol-free medium. mRNA was extracted from cells using an RNeasy Plus Mini Kit (Qiagen), and cDNA was generated using an iScript cDNA Synthesis Kit (Bio-Rad) according to the manufacturer’s instructions. RT-PCR was performed using specific primers for EGR1 and ETV4, and actin was used as a loading control. Data shown are average fold change of I-/I+ ± SD, n = 3 (*, p ≤ 0.05; **, p ≤ 0.01; ***, p < 0.001).

**Table S1.**
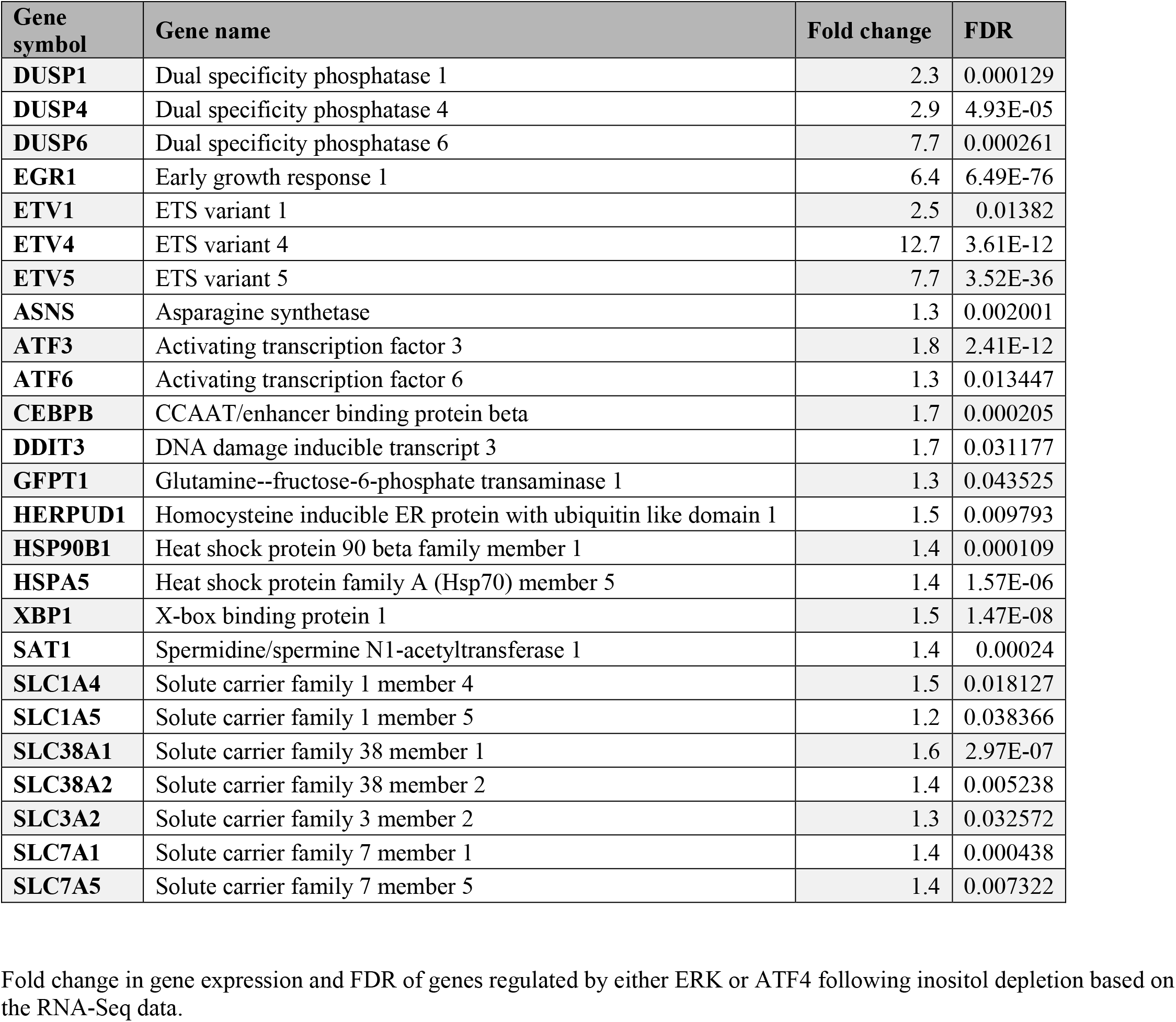
Inositol-sensitive genes regulated by either ERK or ATF4.

### Microsoft Excel files

**Table S2: Lipidomic analysis of HEK293T KO cells following inositol deprivation**

HEK293T ISYNA1-KO cells were cultured for four days in inositol-free medium. Cells were washed with PBS, lysed, then phospholipids extracted and analyzed using HPLC-MS. Data were normalized to the protein content of each sample.

**Table S3: RNA-Seq analysis of HEK293T KO cells following inositol deprivation**

HEK293T ISYNA-KO cells were cultured for four days in inositol-free medium. Total RNA was extracted from cells using TRIzol Reagent (Invitrogen) following the manufacturer’s instructions. Expression analysis was conducted in collaboration with the Wayne State University Genome Sciences Core as described previously.

**Table S4. Off-target analysis of HEK293T WT and ISYNA1-KO cells**

Whole-genome sequencing and differential variant analysis (Psomagen Inc., Rockville, MD) were performed on both the HEK293T WT and ISYNA1-KO cell lines to identify potential off-target mutations.

**Table S5. Regulation of phospholipid metabolism genes by inositol depletion**

Fold change in gene expression and significance of phospholipid metabolism genes based on RNA-Seq data of HEK293T ISYNA1-KO cells following four days of inositol depletion.

